# Imaging progenitor cell differentiation during central nervous system remyelination using an MRI gene reporter

**DOI:** 10.1101/2025.09.10.674209

**Authors:** Eliane E.S. Brechbühl, Myfanwy F.E. Hill, Civia Z. Chen, Hallie F. Gaitsch, Susana Ros, Luca Porcu, Alan J. Wright, Daniel S. Reich, Robin J.M. Franklin, Kevin M. Brindle

## Abstract

Demyelination, the loss of the myelin sheath from around otherwise intact axons, occurs in several diseases, most notably multiple sclerosis (MS). Demyelinated axons that are not remyelinated are vulnerable to irreversible degeneration and therefore therapies that enhance remyelination have been sought. However, there remains a paucity of suitable outcome measures to assess their efficacy. Magnetic Resonance Imaging (MRI) is a non-invasive imaging modality that is used both preclinically and clinically for the assessment of anatomy and tissue function. Here we describe an MRI technique for following the differentiation of oligodendrocyte progenitor cells (OPCs) into oligodendrocytes during the spontaneous regenerative process of remyelination *in vivo*. OPCs were transduced *in situ* with a lentiviral vector expressing an organic anion transporting polypeptide (Oatp1a1) under the control of the differentiation-specific Myelin Basic Protein (MBP) promoter. Oatp1a1 mediates cell uptake of a gadolinium-based MRI contrast agent (Primovist), allowing detection of the cells in T_1_-weighted MR images. Uptake of the contrast agent is restricted to MBP-expressing cells, which is most highly expressed during myelin sheath formation, thereby allowing progenitor-mediated, and potentially oligodendrocyte-mediated, remyelination to be monitored non-invasively *in vivo* using MRI. These findings provide the foundation for the development of direct methods for assessing the efficacy of pro-remyelination therapies.

## Introduction

The pursuit of therapies for the treatment of multiple sclerosis (MS) has historically focused on the suppression of the immune response with the development of disease modifying therapies (DMTs)^1^. Despite their success in reducing the severity and frequency of the relapsing-remitting phase of the disease, they have had little impact on the currently untreatable progressive phase where the underlying gradual loss of axons is in large part due to deficiencies in remyelination. Thus, there has been a focus on treating progressive MS with pro-remyelinating interventions that will restore myelin sheaths and hence protect against axonal loss^2–7^. While cell transplantation-based therapies remain an option, recent efforts have focused on pharmacological promotion of remyelination by endogenous adult oligodendrocyte progenitor cells (OPCs), an abundant type of adult central nervous system (CNS) stem cell. At present, there are several agents with pro-regenerative properties in preclinical studies that are the basis of recent or planned clinical trials, including GPR17^8^, metformin^9^, bexarotene^10^, and clemastine fumarate^11,12^.

Currently, assessing the efficacy of a new myelin regenerative medicine in pre-clinical studies is largely based on histological assessment, and in clinical trials on changes in clinical imaging. However, both these approaches have limitations. In preclinical studies, efficacy assessment usually involves studying oligodendrocyte numbers, and new remyelinated sheaths with a range of microscopy techniques but requires the sacrifice of significant numbers of animals at multiple timepoints. In clinical trials, as in diagnostic imaging, assessment of lesions is made largely using non-invasive imaging techniques, such as MRI, where there is often an imprecise correlation between image appearance and clinical state^13–17^. Additionally, remyelination can be challenging to differentiate from other tissue changes which occur during the resolution of injury including oedema, axonal loss, or cellular infiltration^15,18^. Multiple MRI methods have been developed in order to address this issue including Magnetization Transfer ^19–24^, Diffusion Tensor ^18,25–28^ and Diffusion Kurtosis Imaging^29,30^ and determination of Myelin Water Fraction^31–33^ and a variety of fluorescence^34^ and Positron Emission Tomography techniques^35–38^. However, there remain no imaging modalities for the specific identification of remyelination, unambiguously distinguishing it from other tissue changes.

Here we describe an MRI technique that allows identification of differentiating myelin-forming oligodendrocytes by detecting cells that express high levels of myelin basic protein (MBP). The technique uses an MRI gene reporter under the control of the oligodendrocyte-specific MBP promoter so that MRI contrast is only generated when cells express high levels of MBP, such as when an OPC differentiates into a myelinating oligodendrocyte. The MRI gene reporter is an endogenous transmembrane transporter (Oatp1a1) that, when expressed, facilitates cell uptake of a clinically approved gadolinium-based MRI contrast agent, which generates positive contrast on T_1_-weighted images^39^. Remyelination was imaged in a model of demyelination in which remyelination takes place through the generation of new oligodendrocytes derived from OPCs that had been transduced *in situ* with a lentiviral vector expressing the MBP-Oatp1a1 construct. Contrast is generated in the last stage of remyelination when the OPCs differentiate into new remyelinating oligodendrocytes^40^. The system provides the basis for assessing the efficacy of pro-remyelination therapies for MS and other demyelinating disorders. It is also the first report of assessing stem cell differentiation with MRI and opens up new avenues for exploring stem cell-mediated processes non-invasively.

## Materials and Methods

### Lentiviral production

A PGK (phosphoglycerate kinase) or a rat MBP (myelin basic protein) promoter were used to drive transcription of the red fluorescent protein mStrawberry, and the organic anion transporting polypeptide 1a1 (Slco1a1, NCBI reference sequence NM_017111.1). The mStrawberry coding sequence was separated from the Oatp coding sequence by an E2A sequence, which resulted in equal expression of mStrawberry and Oatp from a single mRNA transcript. The generation of a PGK-mStraw-Oatp plasmid, has been described previously^39^ and the MBP-mStraw-Oatp plasmid was cloned from it by replacing the PGK promoter sequence with the MBP promoter sequence. The transgenes were assembled into the lentiviral vector, pBOBI^41^ (a gift from the Verma laboratory, Salk Institute, La Jolla, USA). The control plasmid PGK-GFP was purchased from Addgene (pLenti PGK GFP Puro (w509-5), plasmid #19070) and MBP-GFP was purchased from VectorBuilder (Chicago, IL, USA) using the MBP promoter sequence shown in Supplementary Table 1. Lentiviral vectors were produced and purified according to published protocols^42,43^. High-concentration virus was produced by ultracentrifugation^43^.

### Animal studies

Sprague Dawley rats were obtained from Charles River (Margate, UK). Only 10-12-week-old females were used. The experiments and analyses were conducted blind. No statistical methods were used to predetermine sample size. The research was conducted under the Animals (Scientific Procedures) Act 1986, Amendment Regulations 2012, following ethical review by the University of Cambridge Animal Welfare and Ethical Review Body (AWERB).

### OPC Isolation

Neonatal OPCs were obtained as described previously^9^. Briefly, the meninges of dissected brains were removed, and the tissue minced and dissociated. The tissue was then gently triturated with pipettes of decreasing diameter to give a cell suspension. The suspension was then centrifuged in a Percoll gradient and sorted by positive magnetic cell separation (MACS) using the A2B5 antibody (Millipore; MAB312). The purified population of OPCs were counted and plated on 24 well plates coated with poly-D-Lysine (Sigma-Aldrich; A-003-E). The OPC medium was changed 24 hours after isolation and after that, half the medium was replaced every other day. Cells were kept at 37°C, in 5% CO_2_ and 5%O_2_. Cell differentiation was induced by replacing, 20 ng/ml b-FGF (Gibco) and 20 ng/ml PDGF (Gibco) in the OPC medium with 40 ng/mL thyroid hormone T_3_ (Sigma). The cells were kept in this medium for 5 days, with a half-media change every other day. In subsequent text, proliferation medium refers to OPC medium with 20 ng/ml bFGF and 20 ng/ml PDGF and differentiation medium to OPC medium with 40 ng/ml thyroid hormone T_3_.

### Lentiviral infection

Lentiviral titres were determined by infecting HEK cells and counting fluorescent cells using flow cytometry. Following isolation, OPCs were allowed to recover in OPC proliferation medium for 24 h before infection at a multiplicity of infection (MOI) of 7 for 24 h before washing with PBS and adding fresh OPC proliferation media.

### Live Cell Imaging

Cells were imaged using an Incucyte^®^ SX5 Live-Cell Analysis Instrument (Sartorius) at 20x magnification, acquiring phase contrast and red and green fluorescence images every 4 h for 84 h. Images were analysed using Incucyte^®^ Base Analysis Software 2022 (Sartorius).

### Flow Cytometry

Flow cytometric analyses used a MacsQuant VYB instrument (Miltenyi), recording mean fluorescence intensity per cell. Analysis was carried out using FlowJo 10.8.1 software, gating for the cells of interest and excluding cell clusters.

### MRI of Cell Pellets

Neonatal OPCs were incubated in 1 mM Primovist (Bayer) in OPC proliferation medium in 6-well plates (max. 2 x 10^6^ cells/well) for 1 h. For inhibition studies, cells were incubated with 1 mM glyburide (Supelco, PHR1287), a pan-Oatp inhibitor^44^, for 5 minutes before adding Primovist. Cells were then washed once in ice-cold PBS, trypsinized with TrypLE (ThermoFisher, cat.no. 12604013), resuspended in OPC medium, centrifuged at 210g for 3 min and then washed twice in ice-cold PBS. Cell pellets were resuspended in 200 µL PBS and transferred to 300 µL tubes and centrifuged on a benchtop microfuge at 2680g for 1 min and then imaged. Images were acquired at 7 T (Agilent, Palo Alto, CA) using a 40 mm diameter Millipede quadrature volume coil^45^ for ^1^H transmit and receive (Agilent). The water proton T_1_ in the pellets was determined using an inversion recovery fast low angle shot (FLASH) sequence, which was modified to include a non-slice-selective adiabatic inversion pulse^46^. T_1_ maps were acquired from a single slice with the following inversion times: 0.01 s, 0.0205 s, 0.042 s, 0.086 s, 0.177 s, 0.362 s, 0.743 s, 1.522 s, 3.121 s and 6.4 s. A T_1_ map was generated using the VnmrJ 3.1.A software (Agilent). The concentration of Primovist in the cell pellet was determined using eqn. 1.

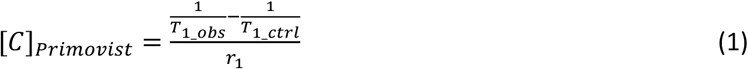

where [C]_Primovist_ is Primovist concentration, T_1_obs_ the T_1_ in the pellet, T_1_ctrl_ the T_1_ in the corresponding control pellet without Primovist added and r_1_ the molar relaxivity of Primovist at 7 T. Images were prepared in FIJI^47^.

### Experimental detail for determining the relaxivity of Primovist

Five serial dilutions of Primovist were prepared in separate small tubes in PBS (0, 0.1, 0.25, 0.5 and 1 mM for the 7T scanner and 0, 0.1, 0.25, 0.4 and 0.5 mM for the 9.4T scanner) and R_1_ maps were acquired with the respective MR scanner. Three replicates were prepared and measured for each tube and the slope on a field-strength specific relaxation rate vs Primovist concentration graph was determined as the relaxivity of Primovist at the respective field strength.

### Calculation of Primovist concentration per cell *in vitro*

c_cell_ = (c_pellet_ * V_pellet_) / N_cells_ Parameters for cell pellet:

- 2 x 10^6^ cells contained in the pellet (N_cells_)

- 2 mm^3^ cell pellet volume (V_pellet_)

- estimated Primovist concentration in the cell pellet (c_pellet_)

### EB-CCP de-remyelination rat model

Bilateral focal demyelinated white matter lesions were created by injecting 4 μl of 0.01% ethidium bromide (EB) into the caudal cerebral peduncles (CCP) of Sprague Dawley female rats and the coordinates of the injection site were recorded^48–50^. Five days post-lesion, 4 μl of high-titre lentivirus were delivered bilaterally into the lesion sites using a stereotaxic frame using the recorded, animal-specific injection-site coordinates. Lentivirus expressing Oatp1a1 was delivered to the left lesion and a control virus to the right lesion. MR images were acquired 13 - 16 days post lesion induction.

### In vivo MRI

Animals were imaged in a 9.4 T spectrometer (Bruker BioSpin) using a ^1^H 72 mm (inner diameter), actively decoupled quadrature birdcage transmit RF coil (Rapid Biomedical) with a ^1^H rat brain actively decoupled receive-only quadrature surface coil (Bruker BioSpin). Intravenous injection of a mannitol solution (0.4 mL, 0.22 µm-filtered 25 % solution) followed immediately by a Primovist solution (3 mL, 50 mM) via a tail vein cannula resulted in an initial serum concentration of Primovist of approximately 8 mM, based on published blood volumes^51^. Animals were anaesthetized using 1.5-3.5 % Isofluorane (Isoflo, Abbott Laboratories, Maidenhead, UK) in air/oxygen (75/25 %, vol/vol, 2 L/min). Body temperature, monitored with a rectal probe, was maintained using warm air at 37°C and breathing rate monitored using a pressure-sensitive pad.

T_2_-weighted images (9 slices, 1 mm thick, repetition time (TR): 1500 ms, echo time (TE): 12.5 ms, 40 x 40 mm^2^ field of view (FOV), 256 x 256 data points, 4 averages per increment) were used to localize the lesions and then a T_1_-weighted image (1 slice, 1 mm thick, TR: 100 ms, TE: 2.5 ms, 40 x 40 mm^2^ FOV, 128 x 128 data points, four averages per increment and flip angle (FA): 35°) of the slice containing the lesions was acquired. An inversion recovery sequence with 8 inversion times (0.02 s, 0.1 s, 0.2 s, 0.5 s, 1 s, 2 s, 3 s, 4 s) was used to produce a R_1_ map of the same slice. The animals were then sacrificed by perfusion fixation with freshly prepared 4% paraformaldehyde (95% purity, Sigma-Aldrich) and their brains dissected for histological examination. Image reconstruction and data analysis were carried out using Matlab (Mathworks, Natick, USA) with a code provided in on GitHub (https://github.com/EliBrec/MRI_Remy). Primovist concentrations within the lesion were calculated using eqn 2.

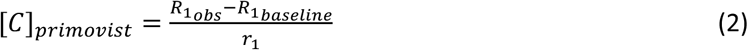

Where C_Primovist_ is the concentration in the EB-CCP lesion, R_1*obs*_ the relaxation rate observed in the lesion at a specific timepoint, R_1*baseline*_ the relaxation rate before Primovist-mannitol co-injection and r_1_ the relaxivity of Primovist at 9.4 T. Images were prepared in FIJI^47^.

### Calculation of Primovist concentration per cell *in vivo*

c_cell_ = (c_lesion_ * V_lesion_) / N_cells_ Parameters for *in vivo* experiments:

- 0.3 x 10^6^ cells contained in lesion (N_cells_)

- 1 mm^3^ lesion volume (V_lesion_)

- estimated Primovist concentration in the lesion (c_lesion_)

### Histology

Brains were post-fixed for 24 h in 4% paraformaldehyde following intra-cardiac perfusion fixation. Tissue was then immersed in 70% ethanol for approximately 5 h before embedding in paraffin and 5 µm sections were cut and placed on glass slides. The sections were then dewaxed (Leica ST5020 multistainer) and rinsed in 95% ethanol before staining in Luxol Fast Blue solution (Luxol Fast Blue Thermo Fisher – 212170250) for 2 h at 60°C. The slides were rinsed in ultrapure water, then in 0.05% lithium carbonate for 5 s and in 70% ethanol for 30-60 s. The sections were then stained with Cellpath Z stain (Mayers) for 5-10 s, washed in ultrapure water for 60 s and 1% eosin for 60 s, rinsed in 70% ethanol and then twice in 100% ethanol, transferred to Xylene for 5 min and a coverslip added (Leica CV5030 coverslipper). The slides were scanned on an Aperio AT2 at 20x magnification with a resolution of 0.5 µm/pixel.

### Quantification of lesion size

Lesion size was determined using the Aperio e-slide viewer by measuring Luxol Fast Blue (LFB)- negative lesions in the caudal cerebellar peduncle (CCP) in the longest direction.

### RNAscope

Detection of rat MBP transcripts was performed on FFPE sections using Advanced Cell Diagnostics (ACD) RNAscope® 2.5 LS Duplex Reagent Kit (Cat No. 322440), RNAscope® 2.5 LS Probe- Mm-Mbp-C2 (Cat No. 451498-C2) (ACD, Hayward, CA, USA). Briefly, 5 µM sections were baked for 1 h at 60°C before loading onto a Bond RX instrument (Leica Biosystems; software version 5.2). Slides were deparaffinized and rehydrated before pre-treatment using Epitope Retrieval Solution 2 (Cat No. AR9640, Leica Biosystems) at 95°C for 15 min, and ACD Enzyme from the Duplex Reagent kit at 40°C for 15 minutes. Probe hybridisation and signal amplification were performed according to manufacturer’s instructions. Fast red detection of rat MBP was performed on the Bond Rx using the Bond Polymer Refine Red Detection Kit (Leica Biosystems, Cat No. DS9390) according to the ACD protocol. Slides were then heated at 60°C for 1 h, dipped in Xylene and mounted using VectaMount Permanent Mounting Medium (Vector Laboratories Burlingame, CA. Cat No. H-5000). The slides were imaged on an Aperio AT2 (Leica Biosystems) to create whole slide images. Images were captured at 40x magnification, with a resolution of 0.25 microns per pixel. Quantification of RNA transcripts per cell was carried out using the HALO image analysis platform (Indica labs).

### Statistics

A first-order delta method was applied for propagation of the standard error in calculations of contrast agent concentration from T_1_ measurements and a multiple comparison z-test with the Holm method was used to adjust the p values when evaluating differences between the cell pellets. *In vivo* data were assessed by a 2-way ANOVA with multiple comparisons.

Statistical analyses were carried out using GraphPad Prism 9, version 9.5.0 (GraphPad software, LLC) or RStudio, version 2023.06.0+421 (Posit software PBC, Boston, MA). p<0.05 was considered statistically significant.

## Results

### Oatp1a1 can be functionally expressed in OPCs

We first showed that OPCs could be transduced with a lentiviral vector to express Oatp1a1. The lentiviral construct contained the sequence for the fluorescent protein mStrawberry linked to the rat Slco1a1 sequence (encoding Oatp1a1) by an E2A sequence, with both under the control of the constitutive PGK promoter (Fig. 1A). Linking via an E2A sequence results in equivalent expression of both transgenes^38^. Following infection of primary neonatal OPC cultures with this PGK-mStrawberry-Oatp1 construct, mStrawberry was detectable within 48 h (Fig. 1C), and transduction efficiency 5 days after infection was 91±3% (n=3) (Fig. 1B). Incubation of OPCs expressing Oatp1a1 with 1 mM Primovist for 1 h and subsequent MR imaging of the cell pellets demonstrated uptake of the MRI contrast agent and therefore expression of functional Oatp1a1 (Fig. 1D). There was a 2.4-fold difference in signal between transduced and non-transduced cells in T_1_-weighted images acquired using an inversion recovery sequence with an inversion time of 1.52 s (Fig. 1E). The mean concentration of Primovist in the pellet was 123 µM (Fig. 1F), as determined from R_1_ (Fig. 1G) and the relaxivity of Primovist at 9.4 T (Suppl. Fig. 1), resulting in approximately 12 x 10^-8^ nmol/cell.

**Figure 1.**
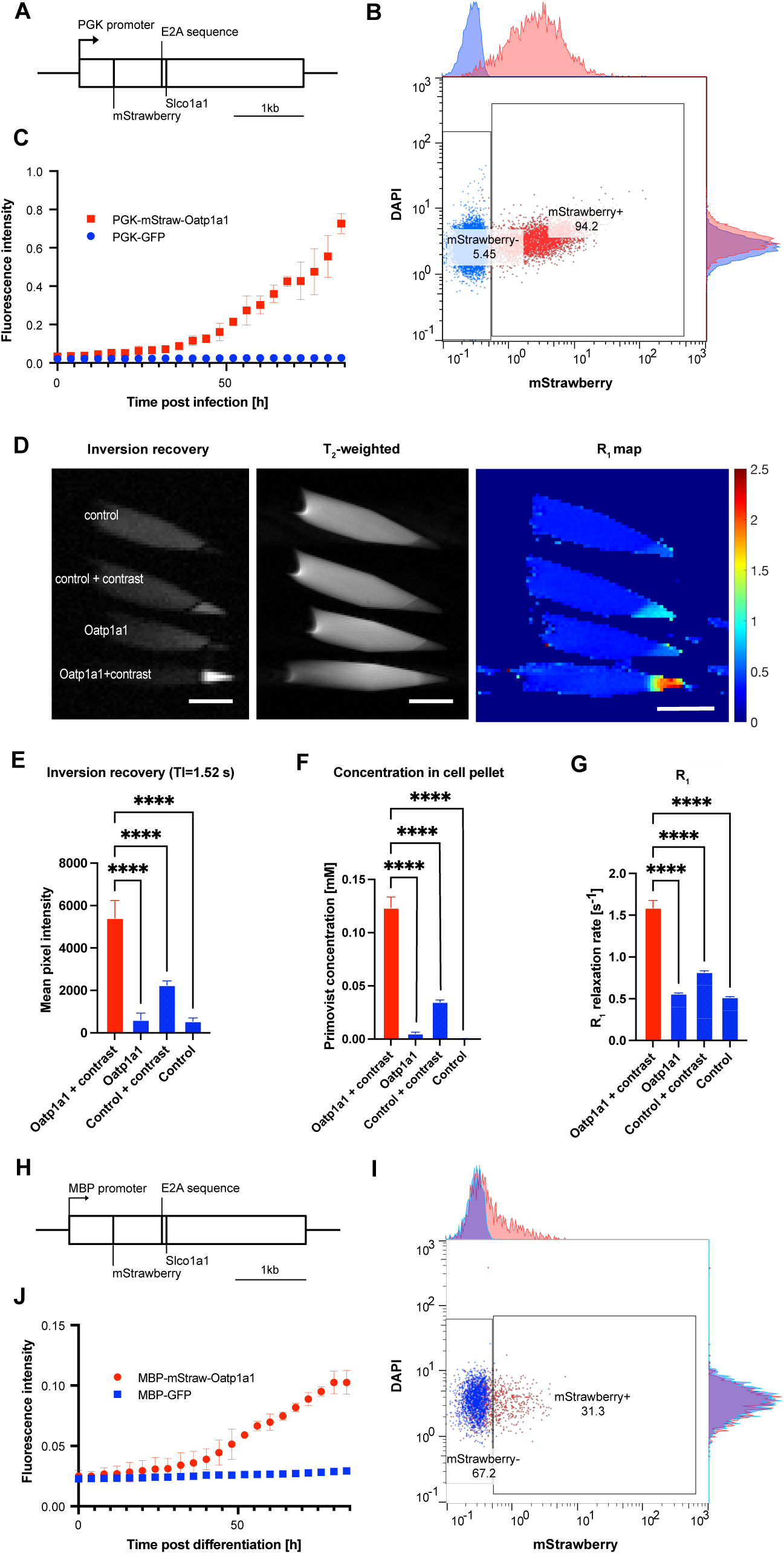
OPCs can be transduced to express Oatp1a1 under the constitutive PGK promoter whereas the MBP promoter results in differentiation-specific expression of Oatp1a1 in OPCs. (A) Plasmid construct used to express Oatp1a1. Oatp expression is controlled by the PGK promoter. (B) Flow cytometry showed that 94.2% of the cells were mStrawberry-positive 48 h post infection. (C) Average mean mStrawberry fluorescence intensity per oligodendrocyte progenitor cell following infection with the PGK-mStrawberry-Oatp1a1 lentivirus or a control virus (PGK-GFP). (D) Inversion recovery image (TI=1.52 s), T_2_-weighted image and R_1_ map of cell pellets of non-transduced OPCs (control) and PGK-mStrawberry-Oatp1a1-transduced OPCs (Oatp1a1) with or without addition of contrast agent. Scale bar: 5 mm. Units on R_1_ map: [s^-1^]. (E) Mean pixel intensities of cell pellets in an inversion recovery image (TI = 1.52 s). (F) Quantification of Primovist concentration in cell pellets based on the R_1_ relaxation rates and the relaxivity of Primovist at 7T. (G) R_1_ relaxation rates in cell pellets quantified from R_1_ maps acquired at 7T. (H) Lentiviral construct MBP-mStrawberry-Oatp1a1. (I) Flow cytometry shows that 31.3% of cells were mStrawberry-positive 48 h after initiation of differentiation. (J) Mean mStrawberry fluorescence intensity per oligodendrocyte progenitor cell after infection and differentiation of the cells with the MBP-mStrawberry-Oatp1a1 lentivirus or a control virus (MBP-GFP).

We then investigated if Oatp1a1 can be expressed when under the control of the differentiation-specific myelin basic protein (MBP) promoter. The plasmid construct used is shown in Fig. 1H. Twenty-four hours after infection of cultures of primary neonatal OPCs with the MBP-mStrawberry-Oatp1 construct, we replaced the growth factors b-FGF and PDGF in the proliferation media with T_3_ in order to differentiate the cells into oligodendrocytes^52^. mStrawberry was detectable within 48 h of differentiation (Fig. 1I) and the transduction efficiency 5 days after the start of differentiation was 28±3% (n=3), as measured by flow cytometry (Fig. 1J). Time courses of the fluorescence intensities for both PGK and MBP driven eGFP expression, plus the mStraw-OATP controls are shown in Suppl. Fig. 2.

### OPC differentiation can be visualised and quantified by MRI in toxin-induced demyelinated lesions

Having shown that Oatp can be used to image differentiating OPCs *in vitro*, we next asked whether differentiating OPCs could also be detected during remyelination following experimentally induced demyelination *in vivo*. First, we determined whether Oatp-expressing cells that had been transduced by intracranial lentiviral administration could take up Primovist delivered systemically (Fig. 2A).

**Figure 2.**
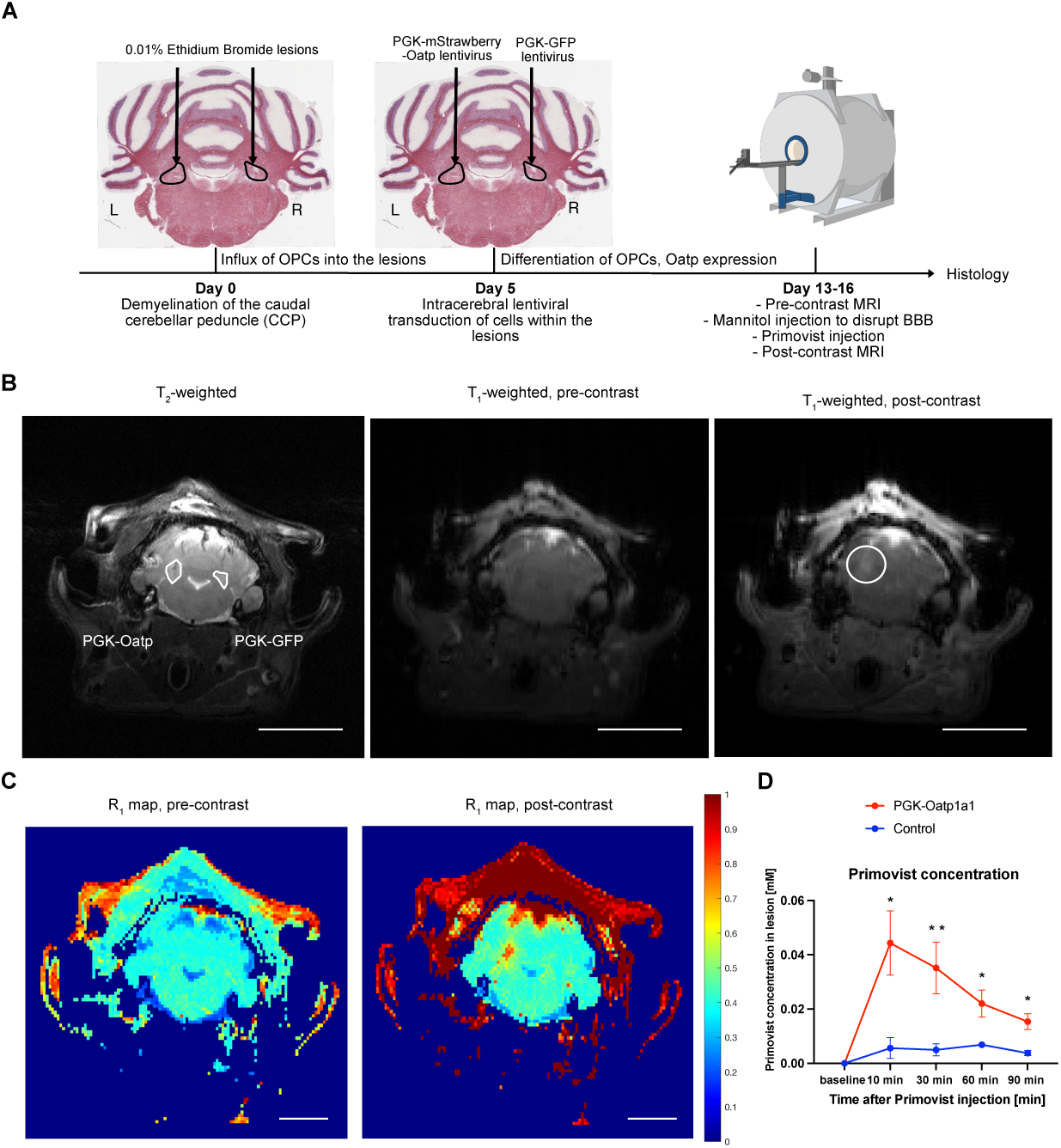
Oatp expression can be detected non-invasively in vivo when under the control of the constitutive PGK promoter. (A) Experimental timeline (B) T_2_- and T_1_-weighted MR images acquired before contrast agent administration and a T_1_-weighted image after Primovist administration. The white circle indicates the lesion containing Oatp-expressing cells. Scale bar: 5 mm Units on R_1_ map: [s^-1^]. (C) R_1_ maps before (left) and 10 min after (right) Primovist administration. Scale bar: 5 mm (D) Quantification of Primovist concentration from r_1_ of Primovist at 9.4 T and R_1_ maps of lesions injected with Oatp-expressing virus and lesions injected with a control virus (PGK-GFP) following contrast agent administration (**p<0.01). (E) Relative signal intensities in inversion recovery images (TI=1 s) of lesions injected with Oatp-expressing virus and lesions injected with a control virus (PGK-GFP) following contrast agent administration (**p<0.01, *p<0.05).

Bilateral caudal cerebellar peduncle (CCP) demyelination was induced using 4 µL ethidium bromide (EB) injections and five days post lesion induction (when endogenous, non-differentiated OPC have populated the lesion^49^), the animals were injected with the PGK-mStrawberry-Oatp lentivirus into the left lesion (4µL containing ͌10^8^ viral particles) and a control virus (PGK-GFP; 4µL containing ͌10^8^ viral particles) into the contralateral lesion. At 13-16 days post lesion induction (when OPCs in the lesion are differentiating into myelinating oligodendrocytes^49^) a T_2_-weighted image was acquired to locate the lesions and to position the imaging slice from which subsequent T_1_-weighted images and R_1_ maps were acquired (Fig. 2B). Following acquisition of a pre-contrast T_1_-weighted image and R_1_ map, a 25% w/v mannitol solution was administered intravenously to open the blood-brain barrier (BBB), followed immediately by administration of Primovist to give an estimated initial blood concentration of approximately 8 mM.

Following Primovist administration, a T_1_-weighted image and an R_1_ map of the same slice were acquired every 20-30 minutes. Representative images of the pre- and post-contrast T_1_-weighted images are shown in Fig. 2B and pre-and post-contrast R_1_ maps in Fig. 2C. Post-contrast, the lesion injected with the PGK-Oatp virus is clearly visible on the T_1_-weighted image and R_1_ map and quantification of contrast agent uptake show that the difference in uptake between lesions injected with the PGK-Oatp virus and with the control virus was significant for up to 30 minutes after contrast administration. The maximum estimated concentration of Primovist in the lesion expressing Oatp was 58 µM and the maximal signal intensity increase (quantified from inversion recovery images at TI=1 s) was more than 5-fold (Fig. 2D and E). After imaging, animals were sacrificed by perfusion fixation and their brains embedded in paraffin for histological examination.

Having shown that endogenous OPCs, amongst other cells, can be transduced with the PGK-Oatp1 lentivirus and imaged *in vivo*, we repeated the experiment using the differentiation-specific MBP-Oatp virus and an MBP-GFP control virus (Fig. 3A). Imaging showed that MBP-driven Oatp expression was detectable on T_1_-weighted images and detectable and quantifiable on R_1_ maps for up to 30 minutes after contrast agent administration (Fig. 3B and 3C). The maximum estimated concentration of Primovist in the lesion expressing Oatp was 15 µM and the maximal signal intensity increase (quantified from inversion recovery images at TI=1 s) was more than 2-fold on T_1_-weighted images (Fig. 3D and 3E).

**Figure 3.**
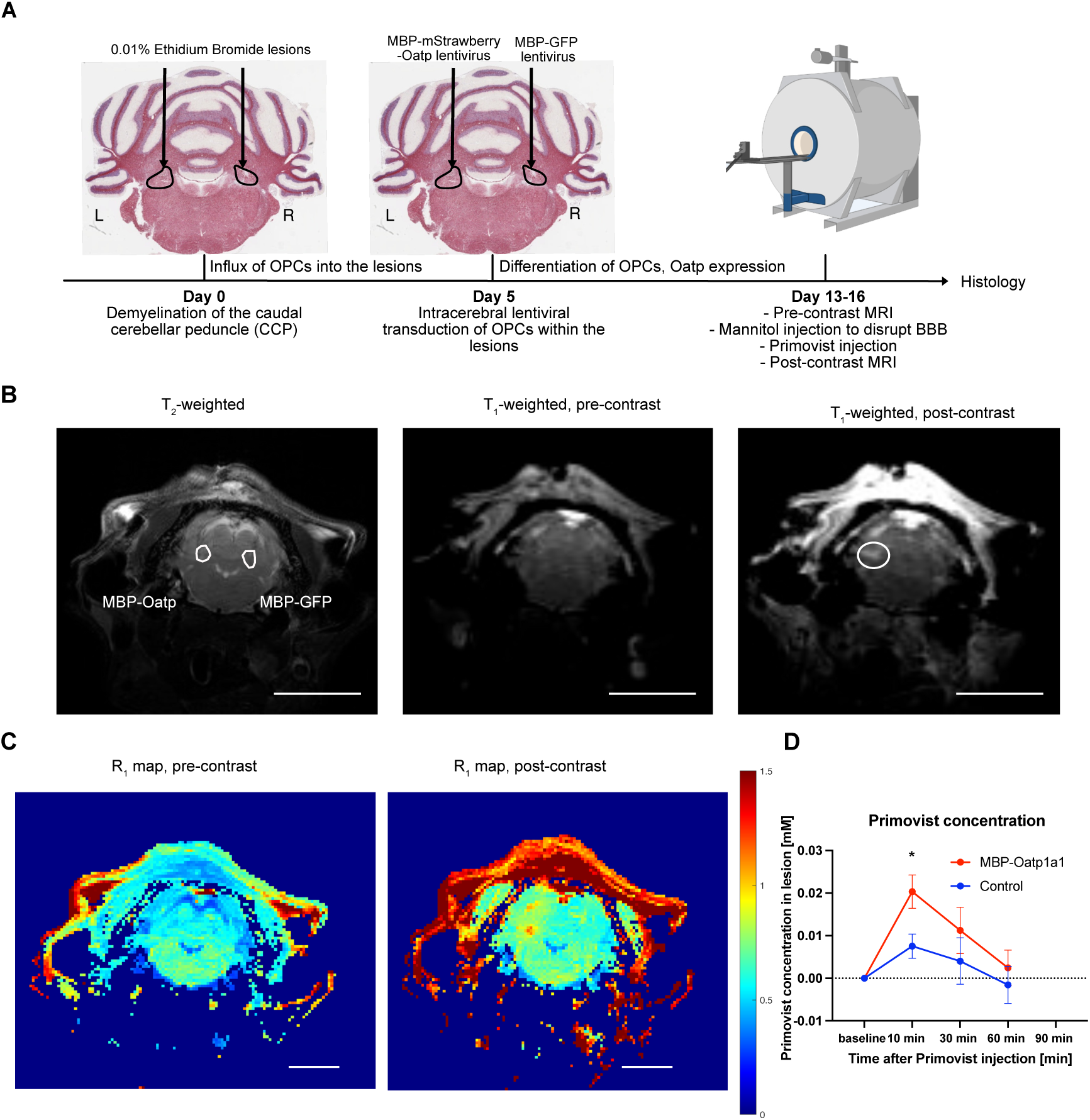
Differentiation-mediated Oatp expression can be detected by T_1_-weighted MRI. (A) Experimental timeline (B) Representative T_2_- and T_1_-weighted MR images before contrast agent administration and a T_1_-weighted image after Primovist administration. The white circle indicates the lesion containing Oatp-expressing cells. Scale bar: 5 mm (C) R_1_ maps pre (left) and 10 minutes post (right) Primovist administration. Scale bar: 5 mm. Units on R_1_ map: [s^-1^] (D) Quantification of Primovist concentration from r_1_ of Primovist at 9.4 T and R_1_ maps of lesions injected with Oatp-expressing virus and lesions injected with a control virus (MBP-GFP) following contrast agent administration (*p<0.05). (E) Relative signal intensities quantified from inversion recovery images (TI=1 s) in lesions injected with Oatp-expressing virus and lesions injected with a control virus (MBP-GFP) following contrast agent administration (**p<0.01).

Histological analysis confirmed that there were no differences in lesion size or cellularity (optical density), or myelin content between lesions containing Oatp-expressing cells and control lesions for both promoters (Suppl. Fig. 3). Further analysis confirmed, by in situ hybridisation, similar levels of mStrawberry and eGFP expression in the lesions into which vector was injected (Suppl. Fig. 4).

Using RNAscope we found that 91% of MBP^+^ cells within and on the rims of the lesion expressed mStrawberry, confirming an excellent transduction rate *in vivo*, which enabled MR imaging (Fig. 4). There was a clear correlation between MBP and Oatp expression, as more than 90% of MBP-expressing cells also expressed mStrawberry, and therefore Oatp, and 73% of mStrawberry-, and therefore Oatp-expressing cells, within the lesion co-express MBP (Fig. 4G). From histological analyses, we counted approximately 3 x 10^5^ Oatp^+^ cells in a lesion of an estimated volume of 1 mm^3^ and from R_1_ maps we calculated a Primovist concentration of 15 µM in a lesion, which results in an estimated intracellular Primovist concentration of 2.5 x 10^-8^ nmol/cell. This estimate of intracellular concentration is on the same order of magnitude as in the *in vitro* uptake experiment (Fig. 1), thus corroborating our findings.

**Figure 4.**
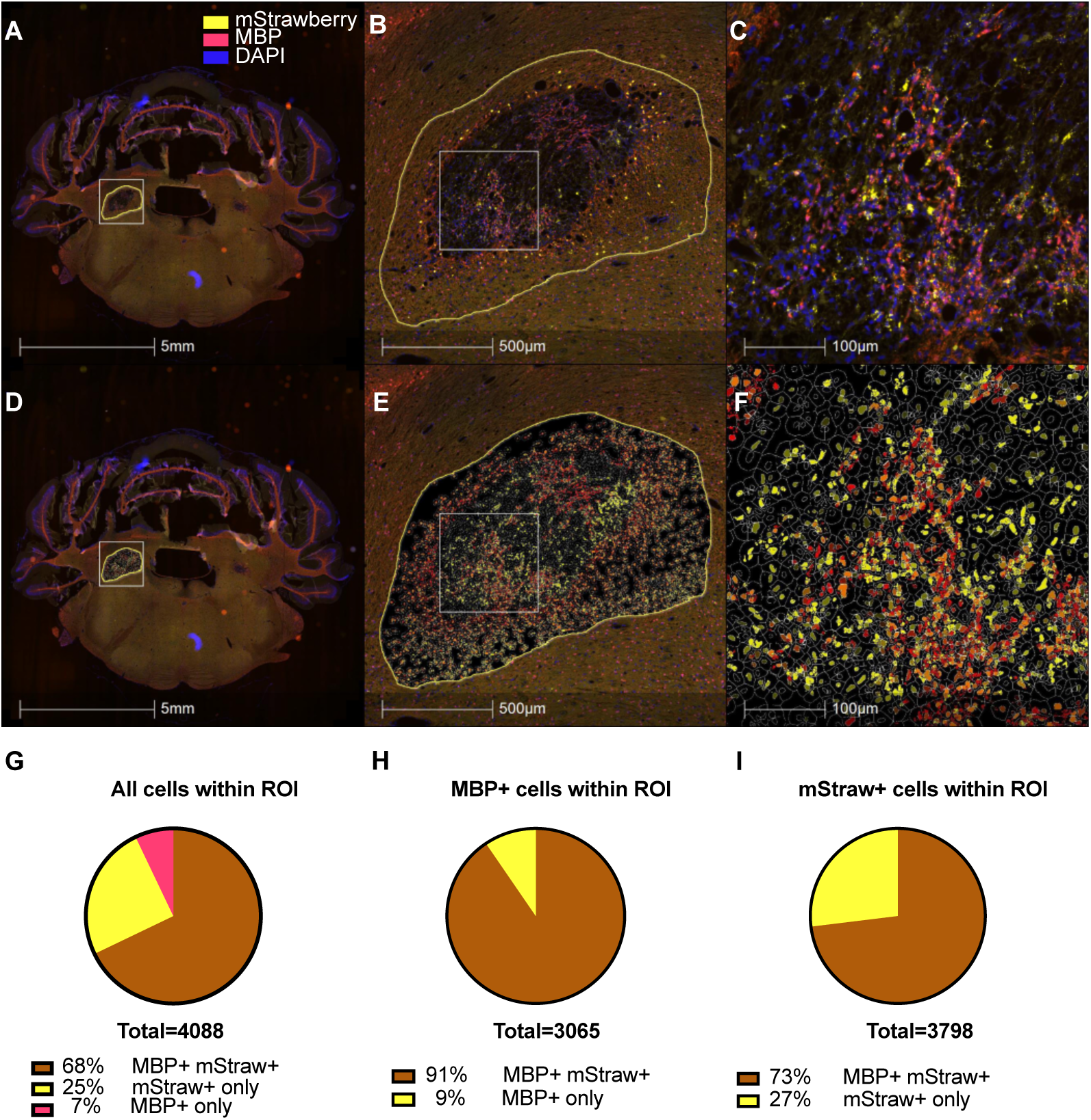
Oatp is preferentially expressed in myelinating cells. (A) Axial section of the cerebellum with a lesion (left) containing Oatp^+^ cells. Part of the control lesion is visible on the contralateral side. (B) Expansion of the lesion from panel A. (C) Expansion of the lesion from panel B. (D) Same panel as A including cell-by-cell analysis showing Oatp^+^ cells in yellow and MBP^+^ cells in red. (E) Same panel as B with cell-by-cell analysis. (G) Same panel as C with cell-by cell analysis. (E) Quantification of percentages of cells within the ROI expressing mStrawberry (readout for Oatp) alone, MBP alone or mStrawberry and MBP together.

## Discussion

Here we have described an MRI gene reporter, Oatp1a1, which, when placed under the control of the MBP promoter, enabled identification of myelin expressing cells in the context of remyelination following experimentally-induced demyelination. Expression of the reporter, which mediates cell uptake of a Gd^3+^-based contrast agent (Primovist), resulted in positive contrast in T_1_-weighted MR images, which is easier to detect than the negative contrast provided by other MRI contrast agents that have been used to track cells, such as Superparamagnetic Iron Oxide Nanoparticles (SPIONs)^53,54^, and other gene reporters, such as ferritin^55^. Isoforms of Oatp have been shown variously to potentiate bioluminescence^56^, and allow detection using NIR fluorescence^57^, photoacoustic^57^ and SPECT^39^ imaging, making Oatp a highly versatile gene reporter for *in vivo* imaging. Moreover, the reversible nature of contrast agent^39^ uptake means that the MBP-Oatp reporter can be used to conduct longitudinal studies of remyelination *in vivo* without the need for tissue collection at multiple time points. The specificity of the differentiation-specific MBP promoter means that cell differentiation and new myelin production can be detected with greater confidence, at an earlier stage, and can be reliably differentiated from several other tissue changes that show similar T_1_-weighted signal changes in MR images, such as oedema^15,18^. With this system, we have been able to image 10^5^ endogenous, primary OPCs *in situ*. This has previously only been possible for cell lines or transplanted primary cells that have been pre-loaded with contrast agents^58^.

To develop an MRI-based reporter of remyelination we used the EB-CCP model^59,60^. This well characterised model has been widely used to study the biology of remyelination, which, while not representing a model of any specific disease, nevertheless allows cellular events and mechanisms involved in remyelination to be explored. CNS remyelination is largely mediated by the generation of new oligodendrocytes from adult OPCs but can occasionally also be mediated by surviving oligodendrocytes within areas of demyelination^61,62^. In the EB-CCP model it is likely that all the remyelination is by oligodendrocytes newly generated from recruited OPCs since demyelination induced by EB arises due to oligodendrocyte death^59^. Thus, in this model, remyelination involves the differentiation of recruited OPCs into new myelin sheath forming oligodendrocytes. This final differentiation phase involves the upregulation of many differentiation-associated genes, including genes that code for key components of the new myelin sheaths. Prominent amongst the latter is the gene for MBP, one of the major protein constituents of the myelin sheath. While MBP continues to be expressed at maintenance levels by steady state myelinating oligodendrocytes, the levels of expression are substantially increased during the differentiation phase during which new myelin sheaths are generated^63^. Thus, although the imaging strategy that we describe here does not directly demonstrate remyelination, it nevertheless provides a powerful surrogate that the process is indeed occurring.

In addition, the gene reporter system described here has translational potential. Primovist is used clinically to image hepatocellular carcinoma^64^; endogenous expression of Oatp in the liver, where it is highly expressed, results in contrast enhancement of liver tissue but not the tumor. Liver uptake is mediated mainly by the OATP1B1 and B3 isoforms and replacement of the rat Oatp1a1 isoform used here with the human OATP1B3 isoform would allow translation of the gene reporter system to the clinic. OATP1B3 has already been used in a gene reporter construct^65^. For example, induced pluripotent stem cells (iPSCs), which are being explored for the treatment of MS ^66,67^, could be transduced *ex vivo* with an MBP-OATP1B3 construct and their differentiation into myelin-forming cells could be imaged *in vivo* using MRI following Primovist administration. Preclinically the MBP-Oatp1a1 reporter could be used to monitor the response of myelinating cells to pro-remyelination therapies^8,9,68,69^, hastening their development and accelerating their clinical translation. More widely, this MRI gene reporter system could be used in the development of other stem cell-mediated regeneration processes.

## Acknowledgements

The authors would like to thank Julia Jones for providing guidance on acquisition and analysis of RNAscope data and the CRUK Cambridge Institute Core Facilities, particularly Caroline Powell in the Histology Core, Richard Grenfell and his team in the Flow Cytometry Core, the Biological Resources Unit, the Research Instrumentation and Cell Services Core Facility and Dominick McIntyre in the Imaging Core.

## Funding

RJMF and DSR acknowledge the Dr Miriam and Sheldon G Adelson Medical Research Foundation, DSR the Intramural Research Program of NINDS/NIH (USA), and KMB and RJMF the UK Multiple Sclerosis Society (MS50) for support. MFEH held a Wellcome Trust doctoral fellowship. HFG was supported by a Gates-Cambridge Scholarship. RJMF was supported by a core support grant from the Wellcome Trust and MRC to the Wellcome Trust-Medical Research Council Cambridge Stem Cell Institute (203151/Z/16/Z). Research in KMB’s lab is supported by Cancer Research UK (C197/A29580, C197/A17242, C9685/A25177).

## Competing interests

The authors declare no conflict of interests with respect to the contents of this paper.

## Supplementary figures

**Supplementary Figure 1.**
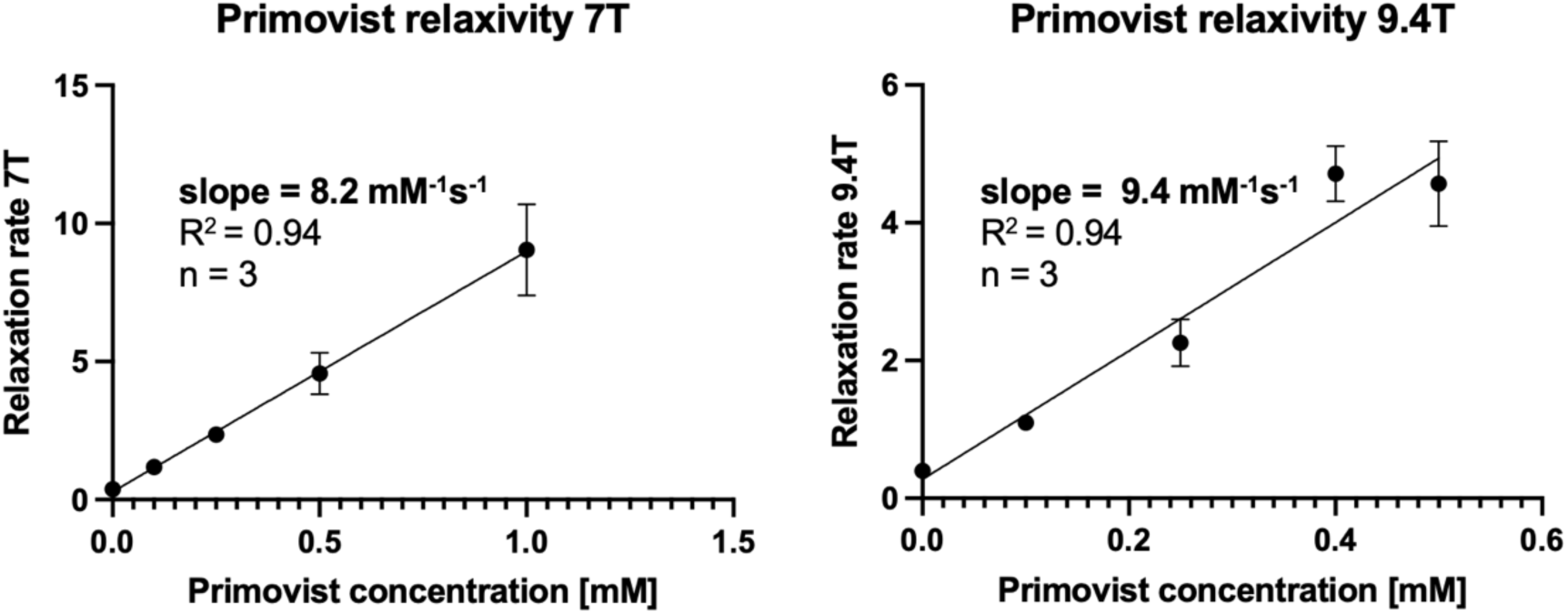
Relaxivity of Primovist at 7 and 9.4 Tesla magnetic field strengths. Five serial dilutions of Primovist were prepared in separate small tubes in PBS (0, 0.1, 0.25, 0.5 and 1 mM for the 7T scanner and 0, 0.1, 0.25, 0.4 and 0.5 mM for the 9.4T scanner) and R_1_ maps were acquired. Three replicates were prepared and measured for each tube and the slope on a field-strength specific relaxation rate vs Primovist concentration graph was determined as the relaxivity of Primovist.

**Supplementary Figure 2.**
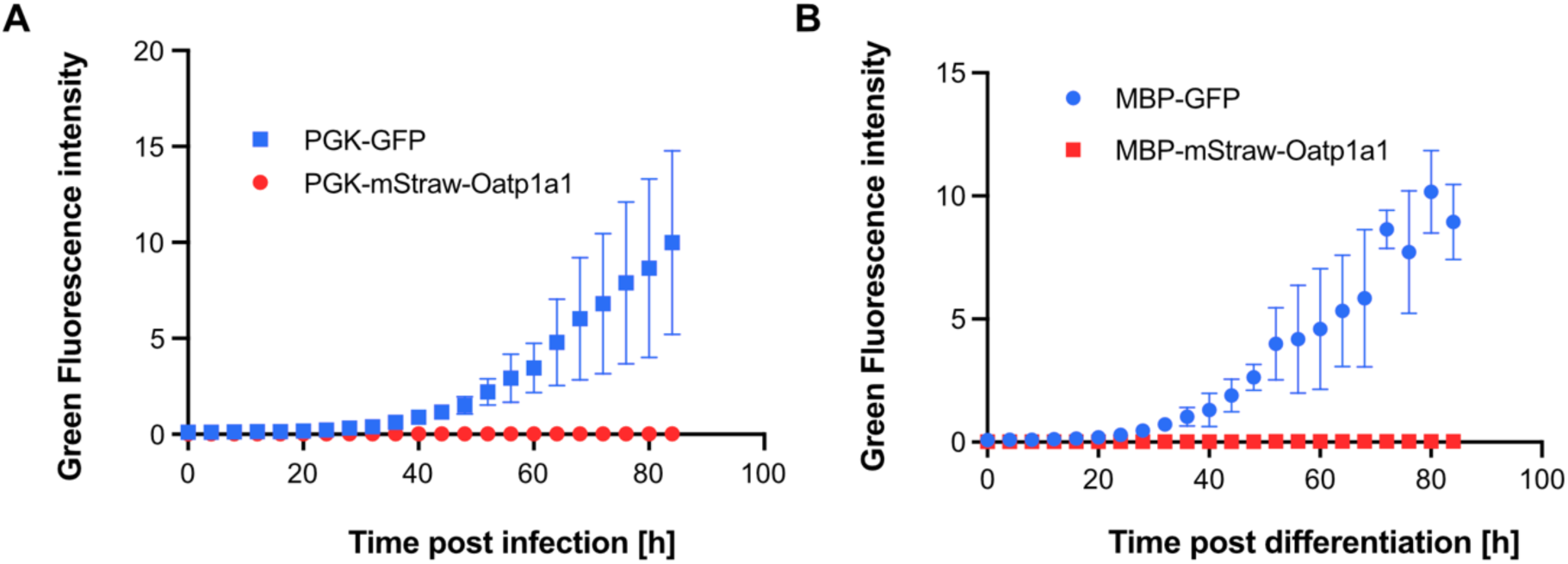
(A) Green Fluorescence Intensity of nOPCs after infection with the control vector PGK-eGFP or PGK-mStrawberry-Oatp. (B) Green Fluorescence Intensity of nOPCs after the start of differentiation, 24 hours after infection with the control vector MBP-eGFP or MBP-mStrawberry-Oatp.

**Supplementary Figure 3.**
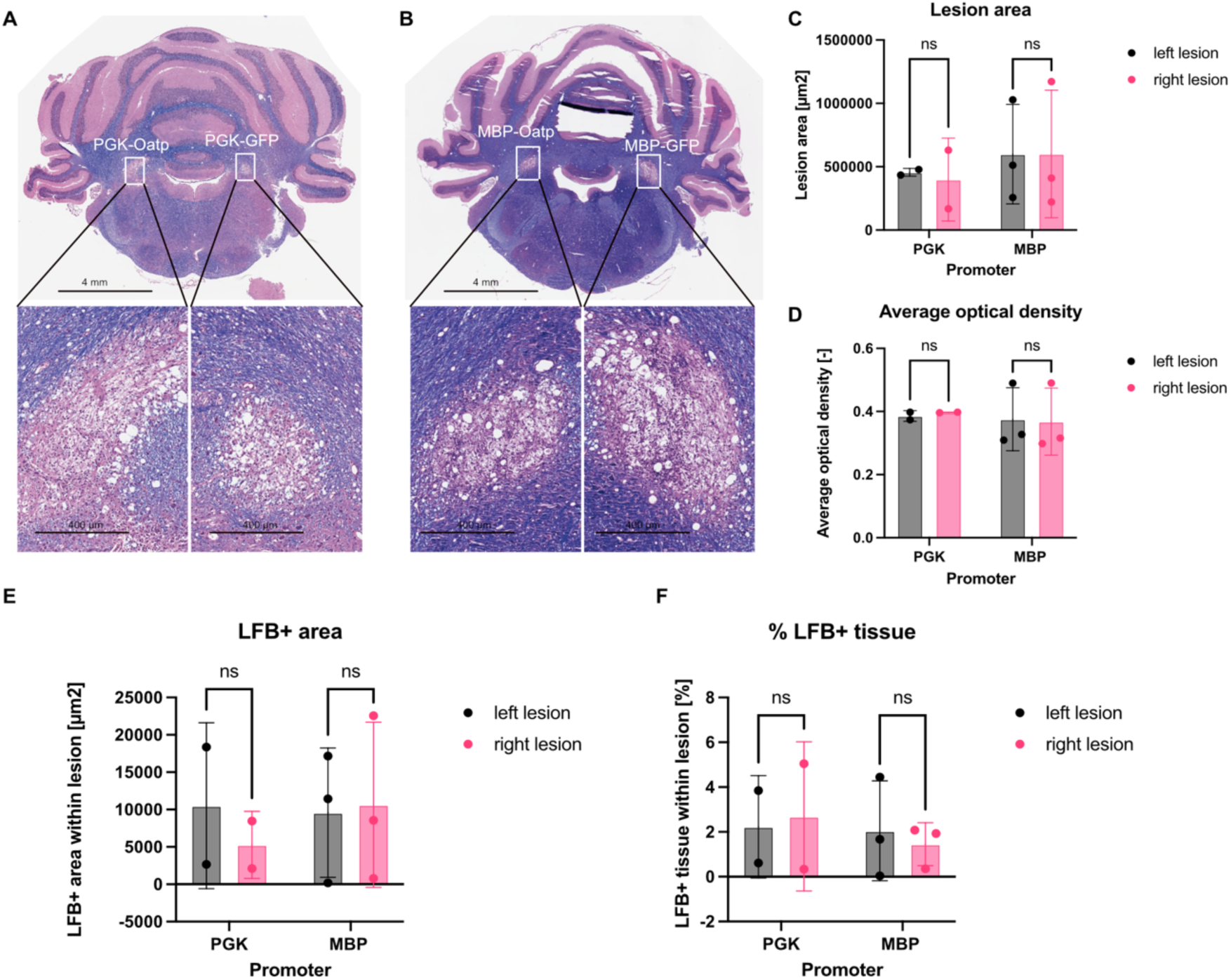
Lesion area, cellularity (optical density) and extent of myelin do not differ between lesions. (A) Representative histological section of cerebellum with bilateral lesions in the caudal cerebellar peduncles (CCP) in which the lesions had been injected with lentiviruses expressing Oatp or GFP under the control of the PGK promoter. A 10X magnification of the lesions show that they had similar cell densities. (B) Representative histological section of cerebellum with bilateral lesions in the caudal cerebellar peduncles (CCP) in which the lesions had been injected with lentiviruses expressing Oatp or GFP under the control of the MBP promoter. A 10X magnification of the lesions show that they had similar cell densities. (C) Lesion area in lesions that had been injected with lentiviruses expressing Oatp or GFP under the control of the PGK and MBP promoter. (D) Average optical density in lesions that had been injected with lentiviruses expressing Oatp or GFP under the control of the PGK and MBP promoter. (E) Area of LFB stain (myelinated area) within lesions for both the experiments using the PGK and the MBP promoter. (F) Percentage of LFB-positive tissue (myelinated tissue) within lesions for both the experiments using the PGK and the MBP promoter.

**Supplementary Figure 4.**
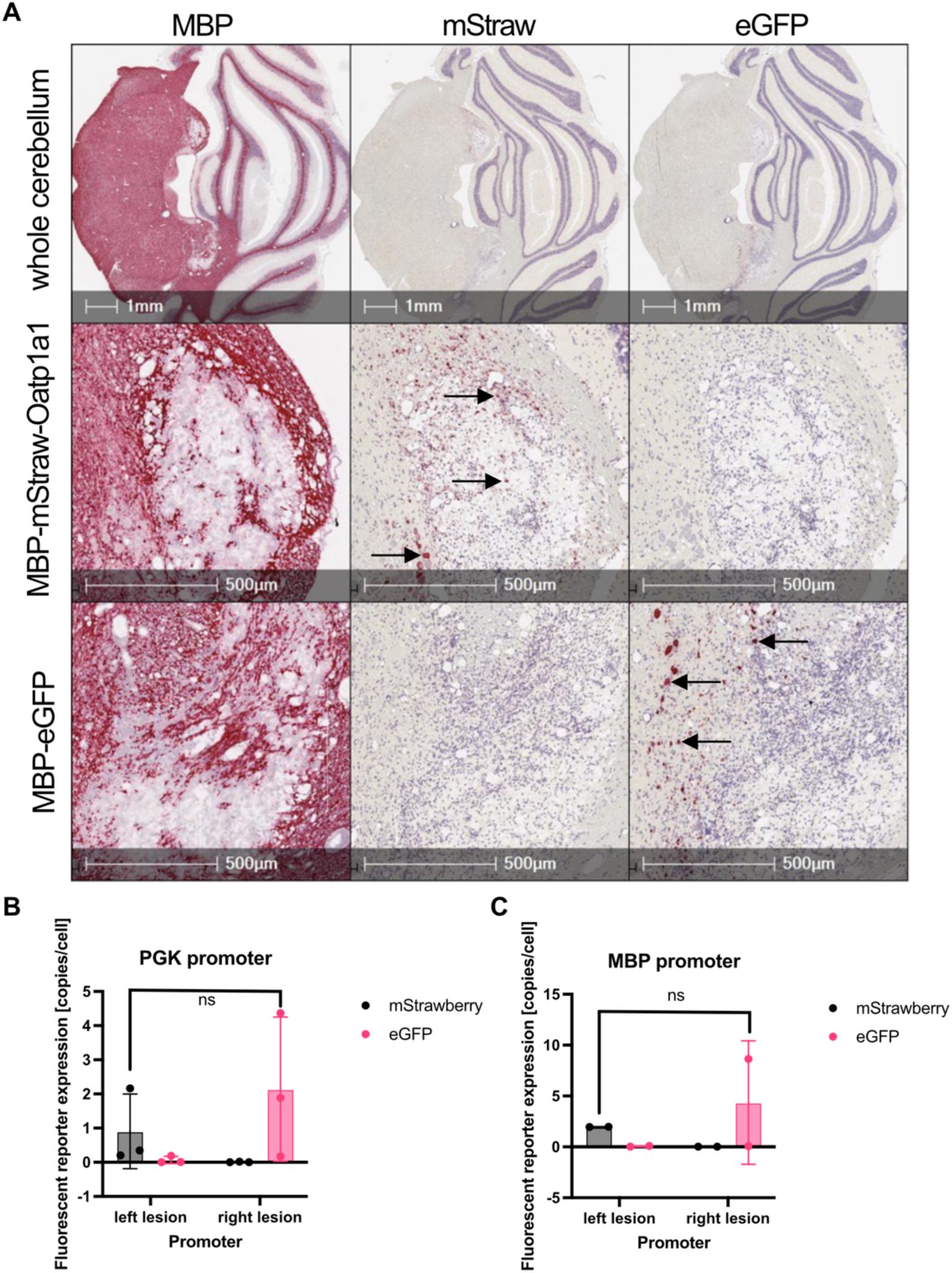
mStrawberry (Oatp) and eGFP are expressed in respective lesions to a similar extent. (A) Top row: Histological sections of the rat cerebellum showing chromogenic *in situ* hybridisation of MBP, mStrawberry and eGFP mRNA. Bilateral lesions in the caudal cerebellar peduncle are visible in the leftmost section (MBP mRNA expression). Middle row: MBP, mStrawberry (mStraw) and eGFP mRNA expression within the left lesion that has received the MBP-mStrawberry-Oatp1a1 virus. mStrawberry mRNA expression can be observed (arrows). Bottom row: MBP, mStrawberry (mStraw) and eGFP mRNA expression within the right lesion that has received the MBP-eGFP control virus. eGFP mRNA expression can be observed (arrows). (B) Expression of mStrawberry and eGFP in both lesions when transgene expression is controlled by the PGK promoter. (C) Expression of mStrawberry and eGFP in both lesions when transgene expression is controlled by the MBP promoter.

## Supplementary table

**Supplementary Table 1.**
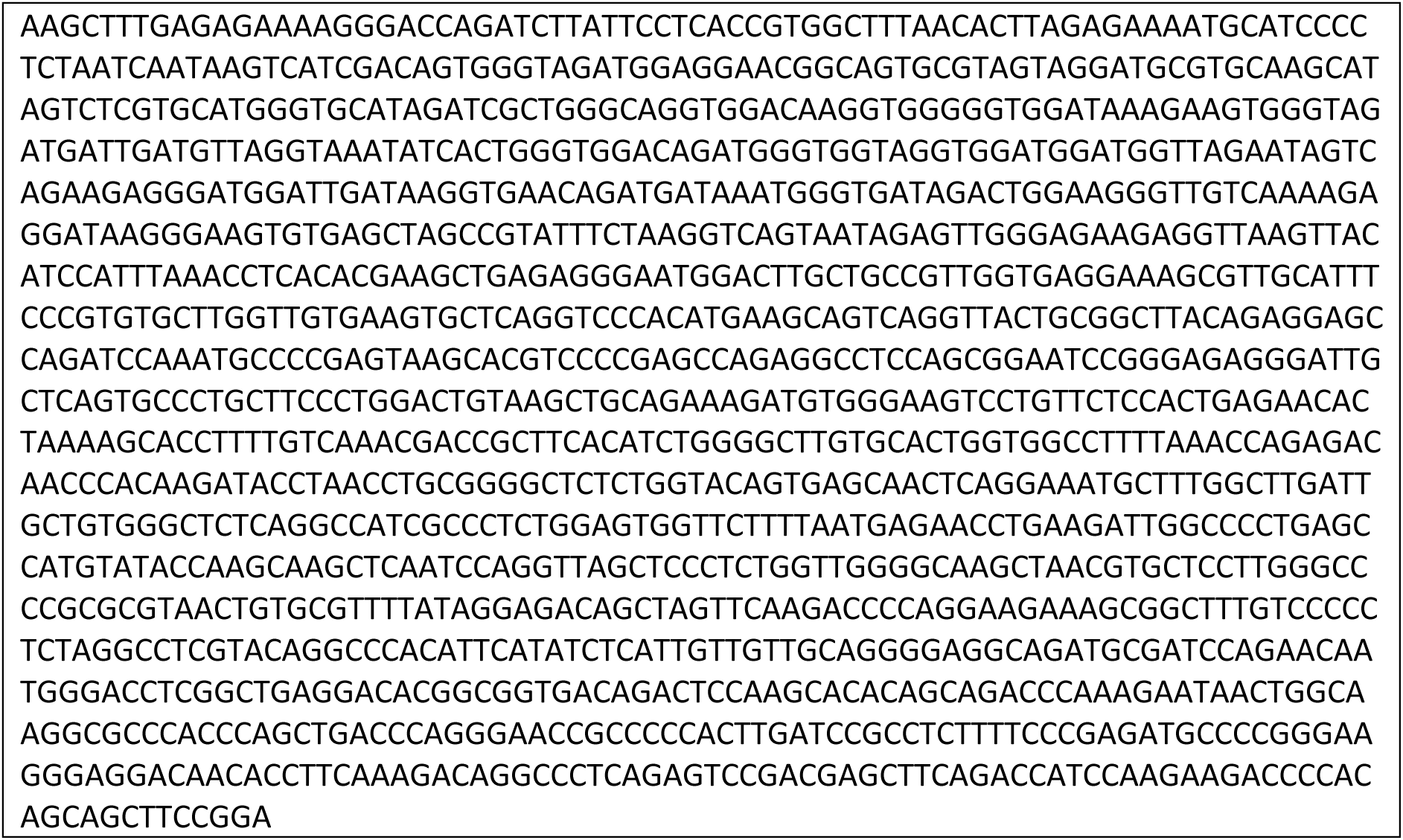
MBP promoter sequence:

## Notes

### Competing Interest Statement

The authors have declared no competing interest.

